# Impaired hepatic autophagy exacerbates xenobiotics induced liver injury

**DOI:** 10.1101/2022.05.27.493749

**Authors:** Katherine Byrnes, Niani Tiaye Bailey, Arissa Mercer, Spandan Joshi, Gang Liu, Xiao-Ming Yin, Bilon Khambu

**Author notes:** **Address corresponding to: Dr. Bilon Khambu**, Department of Pathology and Laboratory Medicine, Tulane University School of Medicine, New Orleans, LA 70112, USA. These two authors share the first authorship (Katherine Byrnes, Niani Tiaye Bailey).

## Abstract

Xenobiotics can activate the hepatic survival pathway, but it is not clear how impaired hepatic survival pathways may affect xenobiotic-induced liver injury. We investigated the role of hepatic autophagy, a cellular survival pathway, in cholestatic liver injury driven by a xenobiotic. Here we demonstrate that DDC diet impaired autophagic flux, resulting in the accumulation of p62-Ub-intrahyaline bodies (IHBs) but not the Mallory Denk-Bodies (MDBs). Impaired autophagic flux was linked to a deregulated hepatic protein-chaperonin system and a significant decline in Rab family proteins. In addition, we demonstrate that heterozygous deletion of Atg7, a key autophagy gene, aggravated the p62-Ub-IHB accumulation and cholestatic liver injury. Moreover, we showed that p62-Ub-IHB accumulation did not activate the proteostasis-related ER stress signaling pathway, but rather activated the NRF2 pathway and suppressed the FXR nuclear receptor, resulting in cholestatic liver injury. *Conclusion*: Impaired autophagy exacerbates xenobiotic-induced cholestatic liver injury. Promotion of autophagy may represent a new therapeutic approach for xenobiotic-induced liver injury.

## INTRODUCTION

Xenobiotics are metabolized and excreted primarily by the liver. Xenobiotics or their reactive metabolites and intermediaries injure tissue and cells, causing liver dysfunction. Host cells respond to xenobiotic exposure by activating survival mechanisms such as macroautophagy, hereafter referred to as autophagy, to minimize the damaging effects of the xenobiotics. Despite extensive research into xenobiotic-mediated cell death (apoptosis and necrosis), little attention has been paid to how host survival mechanisms might mitigate xenobiotic-induced cell death and tissue damage.

Autophagy is an intracellular lysosomal degradative pathway widely known for nutrient turnover, organelle, or signaling protein quality control. In autophagy, cellular proteins and organelles are wrapped in double-membrane vesicles called autophagosomes[1, 2]. The autophagosomes then fuse with the lysosome to form autolysosomes and then use lysosomal degradative enzymes to degrade the cellular contents sequestered in the autophagosomes. A common response to xenobiotic exposure is the formation of misfolded protein aggregates[3]. As a protective mechanism, autophagy helps clear these aggregates, aiding in cellular quality control and ameliorating cellular damage.

DDC (3,5-diethoxycarbonyl-1,4-dihydrocollidine) is the most widely used animal model to study xenobiotic-induced liver injury and fibrosis[4]. Chronic feeding of DDC has long been used to study the formation of Mallory-Denk bodies (MDBs)[5], primary sclerosing cholangitis (PSC)[6], porphyrias[6, 7], chronic cholangiopathies[8], biliary fibrosis[8, 9], and ductular reactions[6, 10].

Using an acute DDC diet model of xenobiotic-induced liver injury, we demonstrate that autophagy impairment causes qualitative accumulation of hepatic proteins such as p62 or Ub resulting in intrahyaline bodies(IHB) formation. The intrahepatic accumulation of p62-Ub containing IHB in the liver activates the NRF2 signaling pathway and downregulates the FXR nuclear receptor, causing bile acid (BA) deregulation. Furthermore, impaired autophagy exacerbates IHB formation and cholestatic liver injury. Thus, autophagy protects against xenobiotic toxicity by regulating the formation of p62-Ub IHBs and modulating p62-NRF2-FXR linked cholestatic liver injury.

## MATERIALS & METHODS

### Animal experiments

All animals used in these experiments were treated with humane care, in accordance with the guidelines from the National Institutes of Health’s Guide for the Care and Use of Laboratory animals, and with approval from Institutional Animal Care and Use Committee (IACUC) of Tulane University. Wild type male mice(C57BL/6J) were randomly assigned to the regular diet fed group and the DDC-diet fed group. Atg7+/-mice((C57BL/6J background) were generated by crossing Alb-Cre mice and the Atg7 F/F control mice as described previously[11].

### DDC diet

Adult mice of the same age(2 month, approximately 25 gram) were randomly assigned to either the regular diet group or the DDC diet group. Those in the regular diet group were fed a regular chow diet of commercially available mouse food pellets (free of xenobiotics) Mice in the DDC diet were fed a diet containing 0.01% of 3,5-diethoxycarbonyl-1,4-dihydrocollidine (DDC) for 2-10 weeks. The mice were then humanely sacrificed, and their livers were collected and frozen to be used for experimental preparations.

### Immunofluorescence microscopy

Paraffin sections were prepared from the liver samples using a microtome. After deparaffinization and rehydration, these sections were treated with a citrate buffer (pH 6.0) for antigen retrieval. The slides were then permeabilized and blocked with 5% donkey or goat serum in PBS with 0.1% Triton X and glycine for one hour. The slides were incubated at 4°C in a solution of primary antibodies diluted in PBS (**Supplementary antibody list Table 1**). The slides were then washed in PBS with 0.1% Triton X and then incubated with a solution of PBS containing fluorophore-conjugated secondary antibodies for at least one hour. Nuclei were stained with Hoechst 33342 (1 μg/ml). Images were obtained using a Nikon Eclipse TE 200 epi-immunofluorescence microscope and the companion NIS-Elements AR3.2 software.

### Serum Parameters analysis

Blood samples were collected and serum analyzzed for ALT, TBA, TB, TC, and TG using commercially available kits from Pointe Scientific, following the manufacturer’s protocol.

### Immunoblotting

Total liver protein lysate was prepared from the liver samples using RIPA lysis buffer contaiing protease inhibitor cocktail.The liver lysate was centrifugation after homogenizing the samples using an electric homogenizer.Sample total protein lysates’ concentrations were measured using a BCA assay (using commercially available kit following standard protocol) and then separated by sodium dodecyl sulfate polyacrylamide gel electrophoresis (SDS-PAGE). After SDS-PAGE, the proteins were transferred to PVDF membrane. The membranes were blocked in TBS with 0.1% Tween 20 (TBS-T) and 5% non-fat dry milk powder for at least one hour. The membranes were then incubated with primary antibodies(**Supplementary antibody list Table 1**) overnight at 4□°C. The membranes were washed with TBS-T and incubated with a horseradish peroxidase-conjugated secondary antibody for 1□hour. Blots were visualized using the immunobilion chemiluminescence system(Millipore, MA) kit and BioRad chemiluminiscence machine scanner.The densitometry analysis of immunoblot images was performed using quantity One Software(Bio-Rad). Densitometry values were normalzed to the loading control(Gapdh or Actin) and then converted to unites relative to the untreated control.

### Quantitative PCR

A GeneJET RNA Purification Kit (Thermo Fisher Scientific) was used to extract the total RNA from homogenized liver samples, according to the manufacturer’s protocol. 1 μg total RNA was then used to synthesize cDNA using a M-MLV Reverse Transcriptase Enzyme System (Life Technologies, Thermo Fisher Scientific) and OligoT primers. qPCR was performed using SYBR Green Master Mixes on a Quanta studio 3 PCR machine (Life Technologies–Applied Biosystems, Thermo Fisher Scientific), using gene-specific primers included in the **Supplementary primer list Table 2**. Gene expression was calculated using the 2^−ΔΔCt^ method and normalized to the housekeeping gene Actin.

### Hematoxylin and Eosin (H&E) Staining

Slides with mouse liver tissue, sectioned in paraffin and fixed in formalin, were stained with H&E according to standard protocol. Images were obtained using Olympus light microscope and analysed using the companion NIS-Elements AR3.2 software.

### Statistical Analysis

Data is presented in figures as the mean with error bars representing ± SEM. SigmaStat 3.5 (Systat Software) was used to perform statistical analysis. Statistical analysis was performed using P values from at least 3 samples, calculated using a 2-tailed Student’s t test for paired group comparisons or 1-way ANOVA with appropriate post hoc analysis for multigroup data comparisons. The statistical analysis methods used were chosen appropriately for variance and distribution fitting. P values of less than 0.05 were considered statistically significant, denoted by “*,” while p values of less than 0.01 were denoted by “**,” and p values of less than 0.001 were denoted by “***.”

## RESULTS

### 1. Acute 3,5-diethoxycarbonyl-1,4-dihydrocollidine (DDC) intoxication leads to accumulation of p62 and Ub containing intracytoplasmic hyaline bodies(IHB) but not Mallory-Denk bodies (MDBs) in liver

The liver synthesizes and secretes millions of protein molecules per day, including plasma proteins[12]. Any abnormality during this process can result in intracellular protein accumulation and cause proteotoxic effects, either through gain or loss of function. Acute feeding of xenobiotic such as DDC, cause hepatic proteinopathy[13, 14]. The development of hepatic proteinopathy is followed by liver injury and various other histological hallmarks of common liver diseases. However, it remains unclear how the DDC diet causes abnormal accumulation of hepatic protein and its mechanism of pathogeneic consequences are less understood.

In order the determine the potential role of cell survival mechanism in DDC mediated liver injury we selected the acute model of DDC intoxication. DDC diet was acutely fed to wild-type mice for 2 weeks to assess how hepatic protein would change. The gross examination revealed enlarged and dark brown livers, which are characteristic of DDC (**Fig.1A**).Interestingly, there was no significant elevation in the level of total hepatic protein level between control and DDC diet-fed mice(**Fig.1B**). The total amount of liver protein in the liver lysate was also not significantly different as determined by the Coomassie Brilliant Blue stain(CBB) (**Fig.1C**). Notably, the protein bands patterns in CBB were quite different in the DDC-exposed liver(**Fig.1C**) indicating a qualitative but not a quantitative change in the hepatic proteome of DDC-exposed liver.

**Figure 1.**
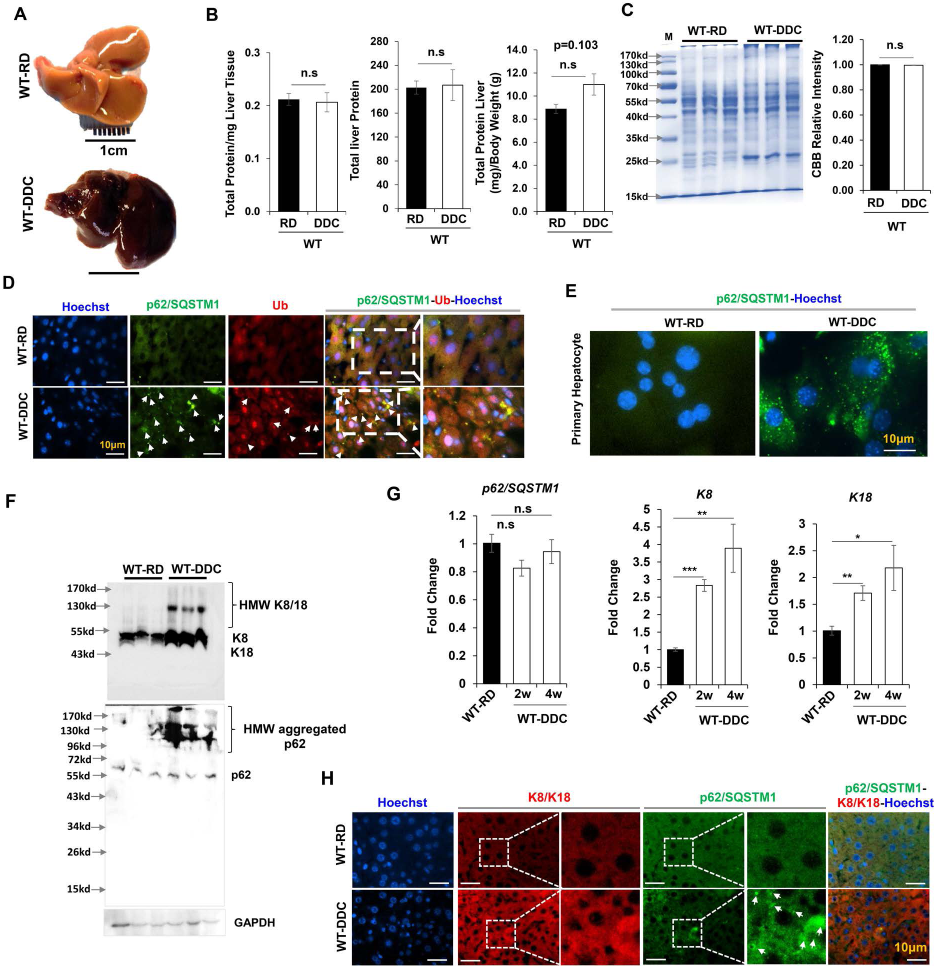
Hepatic proteins are altered qualitatively but not quantitatively by DDC. (A) Gross image of liver harvested from wild type mice fed with regular diet(RD) and 2 weeks of DDC diet. (B)Hepatic total protein level in 9-week-old mice wild type mice fed with RD and 2 weeks of DDC diet. (C) Total liver lysates from 9-week-old mice were analyzed by CBB staining. Protein band intensities were quantified by densitometry.(D) Liver sections were stained for p62/SQSTM1 and Ubiquitin(Ub). Arrows indicate hepatocytes without aggregated p62 or Ub. Scale bars: 10 μm. (E) Immunostained for p62 in primary hepatocytes isolated from 9-week-old wild type mice fed with RD and 2 weeks of DDC diet. (F) Total liver lysates from 9-week-old mice were analyzed by immunoblotting for K8/18, p62, and GAPDH. (G) Quantitative PCR analysis for p62, K8, and K18 mRNA expression in 9-week-old mice wild type mice fed with RD and 2-4 weeks of DDC diet. The mRNA expression levels were normalized to actin. Data are expressed as the mean ± SEM. n.s: not significant,*P≥0.05, **P≥0.01, ***P≥0.001 (n=3).(H) Liver sections were co-stained for K8/K18 and p62. Arrows indicate hepatocytes without aggregated p62. Scale bars:10 μm.

To analyze the qualitative change in the hepatic protein, immunofluorescence staining for p62/Sequestosome-1 was performed. p62/Sequestosome-1 is a major constituent of IHBs[15]. P62 binds Ubiquitin (Ub) and acts as an adapter linking ubiquitinylated species to other proteins. Hence p62 acts as a common denominator in a variety of cytoplasmic inclusions. p62 and Ub proteins, were co-localized and markedly elevated in immunofluorescence stain (**Fig.1D**). A similar accumulation of p62 was noted in primary hepatocytes isolated from mouse feed with a DDC diet for 2 weeks (**Fig.1E**).

Immunoblot analysis also showed marked elevation of p62 and the high molecular weight(HMW) aggregated o62 in DDC diet exposed liver(**Fig.1F**).Notably, mRNA expression did not show any remarkable change for p62(**Fig.1G**). p62 and Ub aggregates can cross-link with the intermediate filament proteins keratins 8 and 18(K8/K18) to form MDBs[14]. The expression levels of K8 and K18, were elevated at both mRNA and protein levels (**Fig.1F-1G**). Increased levels of both normal and high-molecular-weight (HMW) size of K8 and K18 were observed in DDC-exposed liver (**Fig.1F-1G**).

We next examined whether the p62-Ub containing IHBs contains the K8/K18 proteins as IHBs formation generally leads to MDB formation[14] and there was a significant elevation of K8/K18 expression in the DDC diet treated condition(**Fig.1G**). Immunofluorescence staining of K8/K18 did not show the presence of protein aggregates even though immunoblot showed an elevated level of HMW K8/K18(**Fig.1H, Supplementary Fig1A-1B**). More importantly, the K8/K18 did not co-localize with the p62 aggregates(**Fig.1H, Supplementary Fig1A-1B**) suggesting that acute DDC cause protein aggregate of p62 but not the K8/K18.Thus, acute DDC causes a qualitative change in hepatic proteome and likely sets the stage for aberrant cellular stress signaling. Acute DDC induces the formation of a p62-Ub inclusion bodies accumulation that are devoid of cytokeratins, suggesting it can be characterized as IHBs and not MDBs.

### 2. DDC deregulates the hepatic protein chaperone system

Acute DDC exposed liver has an accumulation of p62-Ub IHBs(**Fig.1**).The selective formation of the hepatic IHBs could be due to failure in the molecular chaperone. Heat shock proteins(HSPs) serve as molecular chaperones which guide conformations critical for protein synthesis, folding, translocation, and assembly during cellular stress[16, 17]. During cellular stress, small HSPs, such as HSP70/90 keep the newly formed misfolded proteins in a folding-competent state until the physiological situation is improved. Hsp70 can also allows ATP-dependent refolding of the misfolded proteins. Molecular chaperones such as Hsp25 and Hsp70 were previously described as a component of MDBs[18]. Protein misfolding and subsequent aggregation in various organelles leads to proteotoxicity.Given the impaired proteostasis and protein inclusion bodies (IHBs) observed in the acute DDC-diet model, we hypothesized that deregulated HSPs contribute to improper protein folding, leading to protein accumulation and then proteotoxic liver injury. We thus examined the impact of DCC-diet on the expression of molecular chaperonins.

HSPs belong to seven major families including HSP110, HSP90, HSP70(A), HSP60(D), HSP40(DnaJ), HSP47, and HSP10(E)[19].Of 26 different HSPs expression levels analyzed by qPCR, we found decreased expression of nearly all of the HSPs, across all of the major heat shock families in the DDC-diet fed liver(**Fig.2A-2G**). Distinct members of the HSPs family localize to the cytosol, mitochondria, and endoplasmic reticulum(ER)[20]. Irrespective of the cellular location, all HSPs were downregulated(**Supplementary Fig.2-3**). The downregulation of mRNA of HSPs was properly reflected in their proteins level(**Fig.2H**).

**Figure 2.**
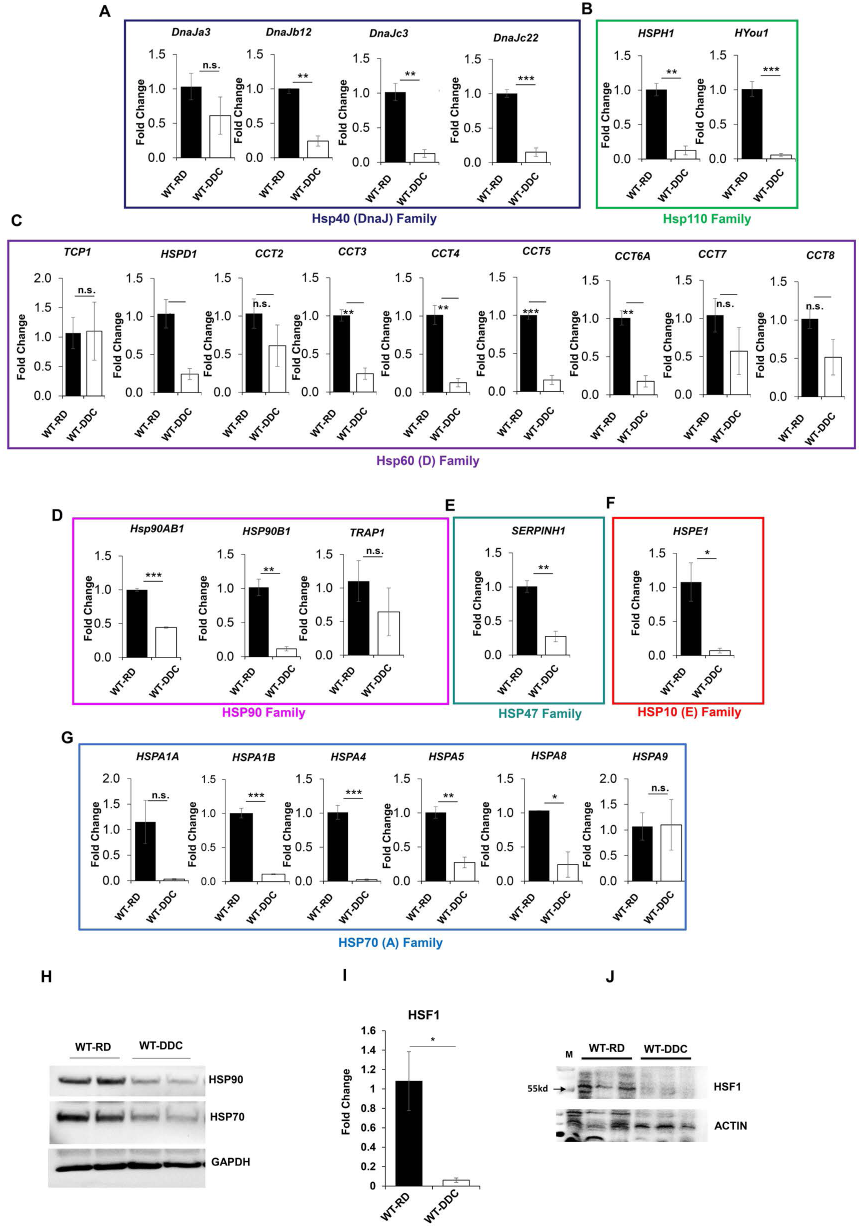
Liver protein chaperonin system is suppressed by DDC. Quantitative PCR analysis for (A) Hsp40 (DnaJ) Family, (B) Hsp110 Family, (C) Hsp60 Family, (D) HSP90 Family, (E) HSP47 Family, (F) HSP10 (E) Family, (F) HSP70 Family, and (G) HSF1 mRNA expression in 9-week-old mice wild type mice fed with regular diet(RD) and 2 weeks of DDC diet. The mRNA expression levels were normalized to actin. Data are expressed as the mean ± SEM. n.s: not significant,*P≥0.05, **P≥0.01, ***P≥0.001 (n=3).(I-J) Total liver lysates from 9-week-old mice fed with RD and 2 weeks of DDC diet were analyzed by immunoblotting for HSF1 or ACTIN(I) and HSP90, HSP70, and GAPDH.

Transcriptional activation of HSPs is orchestrated by heat shock factor 1(HSF1), wchi rapidly translocates to HSP genes and induce their expression[21].So we hypothesized that deregulation of the HSF1 signaling directly contributes to suppression of HSPs in DDC exposed liver.

Notably, the mRNA or protein levels of HSF1 were also suppressed by DDC(**Fig.2I-2J, Supplementary Fig.3**) suggesting that there is ongoing deregulation of the proteostasis signaling pathway in DDC diet-fed liver. Interestingly, refeeding regular diet to DDC fed mice recovered the expression of the HSF1 and HSPs(**Supplementary Fig.4)** suggesting that the suppression of molecular chaperinins and HSF1 is temporariy and reversible.Thus acute DDC can suppress the protein molecular chaperone system, and contribute to the formation of p62-Ub-IHBs. Furthermore, the downregulation of these proteostasis factors could be one possible contributing factor to selective protein accumulation.

### 3. Autophagy flux is impaired by an acute DDC diet

Hepatic IHBs formation could also be due to impaired clearance of the protein aggregates such as by autophagy or ubiquitin-proteasome degradation pathways[22-24]. Hepatocytes have active autophagy even at basal level and can utilize to counteract the proteotoxicity. Moreover, p62 protein which is one of the selective autophagic substrates, accumulates in the DDC exposed liver(**Fig.1D-1F**), Whether DDC diet-mediated IHBs formation is due to impaired hepatic autophagy is less clear. So, we next determined the autophagy status in DDC or regular diet fed mice by examining the level of autophagy-specific marker-LC3-I and LC3-II. Immunofluorescence staining for LC3 protein showed increased LC3 positive autophagic puntaes in DDC fed mice(**Fig.3A, Supplementary Fig.5A**). Western blot analysis also showed the elevated hepatic LC3-II protein levels in DDC fed liver compared to their regular diet fed wild type (**Supplementary Fig.5B**). Examination of p62, an autophagic substrate by immunofluorescence staining showed co-localization of LC3 with p62 autophagic substrate(**Fig.3A**), sugggeting that hepatic autophagic function is altered with the DDC diet.

**Figure 3.**
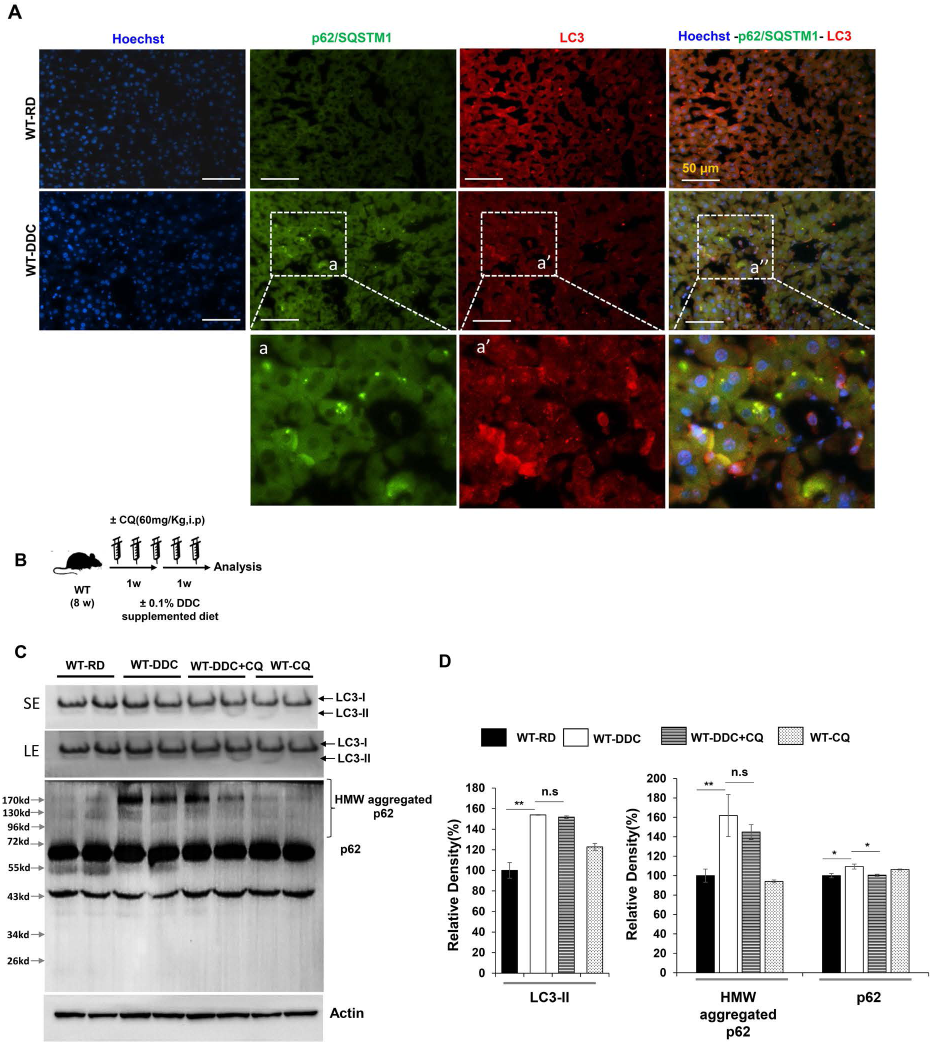
Autophagy flux is impaired by DDC. (A) Co-immunostained for p62 and LC3 in liver section prepared from wild type mice fed with regular diet(RD) and 2 weeks of DDC diet. The a, a’, and a’’ represents the enlarged images from the respective images from DDC diet fed wild type liver. Scale bars: 50 μm. (B) Schematics of CQ administration to Regular diet(RD) or 2 weeks of DDC diet fed wild type mice. (C) Total liver lysates were analyzed by immunoblotting for LC3, p62 and ACTIN. (D) Protein band intensities for LC3-II, high molecular weight(HMW) p62, and normal molecular weight p62 were quantified by densitometry. Protein band intensity was normalized to ACTIN band intensity. Data are expressed as the mean ± SEM. n.s: not significant,*P≥0.05, **P≥0.01 (n=2).

Increased hepatic LC3-II protein level and elevated intracellular autophagic puntaes could be either due to autophagy induction or due to impaired fusion of autophagosomes with the lysosomes[25]. To rule out these possibilities, we performed autophagic-flux in DDC-fed mice. The DDC diet-fed mice were injected with chloroquine(CQ), and an autophagic flux inhibitor(**Fig.3B**). Immunoblotting showed increased LC3-II level in DDC exposed liver but there were no further increase in the LC3-II protein level in presence of CQ(**Fig.3C-3D**).Furthermore the extent of elevation of p62 (HMW or normal molecular weight) levels were similar between CQ treated or not treated DDC exposed liver(**Fig.3C-3D**). These results indicate that that increased autophagosome punctae is not because of autophagy induction rather due to DDC mediated impairment of fusion of autophagosomes with the lysosomes. Overall these data indicate that hepatic IHBs formation (protein accumulation) is not only due to the downregulation of the HSF1 and HSPs signaling network but also due to the impaired clearance of IHBs by autophagy. Whether and how these two cellular events are mechanistically linked or not is unclear.

### 4. Impaired autophagic flux is not due to endoplasmic reticulum (ER) stress signaling

The DDC diet causes impaired hepatic autophagy by blocking the fusion of autophagosomes with the lysosomes and preventing the degradation of p62-Ub IHBs(**Fig.3A-3D**).Autophagic flux impairment is observed during xenobiotic stress such as during thapsigargin treatment and high fat-fed conditions [26-28]. Thapsigargin treatment induces ER stress by inhibiting sarcoendoplasmic reticulum calcium transport ATPase (SERCA) and blocks autophagosome-lysosome fusion[26]. Similarly, free fatty acid (FFA) from a high-fat diet induces ER stress, inhibits SERCA activity-increasing cytosolic calcium in hepatocytes that lead to the inhibition of autophagic function primarily by blocking the fusion step[28]. ER stress could also be activated in response to protein misfolding and aggregate formation in the DDC diet[3].

So we next asked whether the DDC induces ER stress and ER stress-mediated elevation of cytosolic Ca2+ responsible for the hepatic autophagic flux impairment. We examined the status of various ER stress markers in the DDC diet exposed liver. Misfolded protein accumulation leads to the activation of three constitutive client proteins PKR-like endoplasmic reticulum kinase (PERK), inositol requiring enzyme-1 (IRE1), and activating transcription factor 6 (ATF6)) that serve as upstream activators of co-ordinated signal transduction known as unfolded protein response(UPR)[29]. Examination of various signaling proteins concerning the PERK pathway(eIF2a, CHOP, GADD34), ATF6 pathway(ATG6-p90, ATG6-p50, and IREa pathway (XBP-spliced, XBP-unspliced, IREFα) did not show any remarkable upregulation of these proteins level in DDC exposed liver lysate. Instead, expression level of these proteins were downregulated(**Fig.4A**). The mRNA expression level of XBP or chop were also significantly downregulated in DDC exposed liver(**Fig.4B**). Interestingly the suppression of these ER stress proteins appears reversible as the refeeding regular diet after 2-weeks of DDC exposure elevated their mRNA expression than the basal expression level(**Fig.4B**). Thus, ER stress is not activated but rather suppressed by the DDC diet. These results suggests that ER-stress is unlikely to mechanistically explain the observed autophagicflux impairment in the DDC exposed liver.

**Figure 4.**
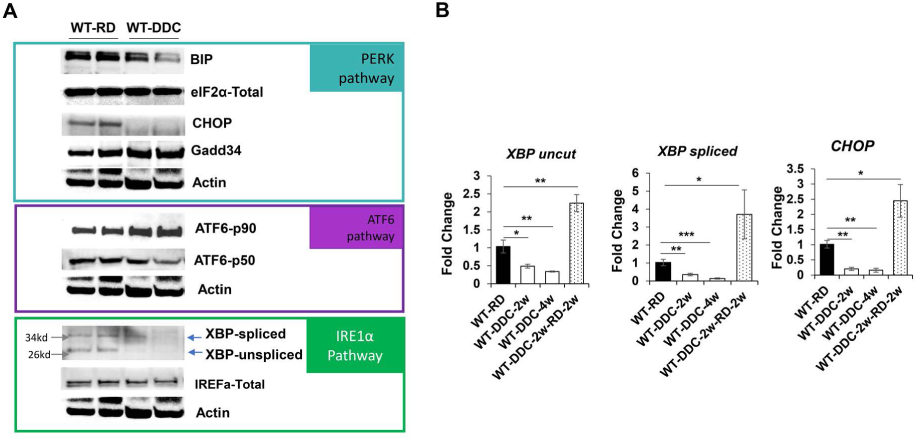
Inhibition of autophagic flux by DDC diet is not triggered by ER stress. (A) Total liver lysates were analyzed by immunoblotting for PERK Pathway(BIP, eIF2a, CHOP, GADD34, ACTIN), ATF6 pathway(ATF-p90, ATF-p50, ACTIN), and (IRF1a pathway(XBP-spliced, XBP-unspliced, ACTIN) proteins. (B) Quantitative PCR analysis for XBP-uncut, XBP spliced, and CHOP mRNA expression from wild type mice fed with regular diet(RD), 2-4 weeks of DDC diet, and 2week DDC followed by RD. The mRNA expression levels were normalized to actin. Data are expressed as the mean ± SEM. *P≥0.05, **P≥0.01, ***P≥0.001(n=3).

### 5. Impaired autophagic flux is correlated to the downregulation of Rab proteins

Autophagic degradation involves the vesicular formation, transport, tethring, and fusion[30]. Autophagic flux particularly involved the fusion of autophagosomes with the lysosome to degrade the enclosed contents including protein aggregates. The Rab protein is a small GTPase that belongs to the Ras-like GTPase superfamily and shuttles between GTP-bound active state and GDP-bound inactive state to have central role in vesicular trafficking[31]. Since autophagy involves series of vesicular trafficking events, Rab small GTPases have been found to participate in the autophagic process[32]. Numerous Rab proteins such as Rab1, Rab5, Rab7, rab9A, Rab11, Rab23, Rab32, and Rab33B participates in autophagosomes[30]. The Rab7 has been particularly shown to be involved in the fusion of autophagosomes to lysosomes[33-36]. So, we next examined whether deregulation of Rab GTPases could be related to impaired autophagic flux in DDC exposed liver. We determined the expression level of the Rab genes in the DDC diet-fed liver and compared to regular diet-fed liver. We identified 59 Rab genes expressed in rodents at a detectable level and selected 36 of these Rabs for quantitative PCR analysis. We particularly focused on the Rabs which are known to be either expressed in the liver or implicated in autophagy. Gene expression analysis showed a general downregulation of Rab genes in DDC diet exposed liver(**Fig.5A**). We verified this observation by examining the protein expression level of a few selected Rabs and Rab-associated proteins such as Rabex5,

**Figure 5.**
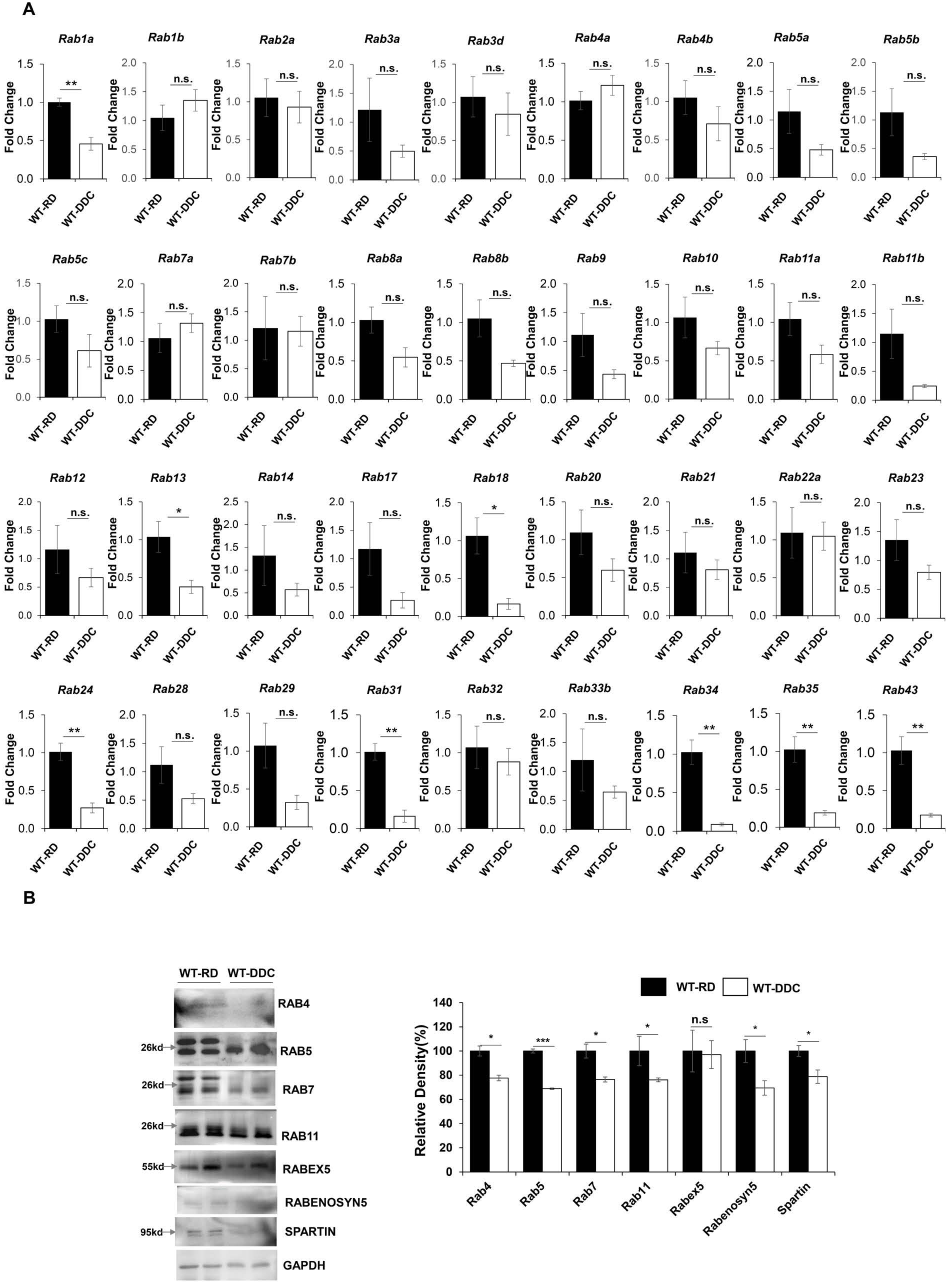
Inhibition of autophagic flux by DDC diet may be due to the downregulation of Rab proteins. (A) Quantitative PCR analysis for Rab family genes in mRNA expression in wild type mice fed with regular diet(RD), and 2 weeks of DDC diet. The mRNA expression levels were normalized to actin. Data are expressed as the mean ± SEM. n.s: not significant, *P≥0.05, **P≥0.01, ***P≥0.001(n=3). (B) Total liver lysates were analyzed by immunoblotting for RAB4, RAB5, RAB7, RAB11, RABEX5, RABENOSYN5, SPARTIN and GAPDH proteins. Protein band intensity was normalized to GAPDH band intensity. Data are expressed as the mean ± SEM. n.s: not significant,*P≥0.05, ***P≥0.001 (n=2).

Rabenosyn5, and Spartin. Immunoblotting analysis showed significant downregulation of protein levels of Rab4, Rab5, Rab7, Rab11, Rabex5, Rabenosyn5, and Spartin in the DDC diet exposed liver(**Fig.5B-5C**). These data suggest that suppression of Rab GTPase expression and function impairs autophagic flux and causes qualitative changes in the hepatic proteome of DDC exposed liver.

### 6. Impaired hepatic autophagy exacerbates DDC induced cholestatic liver injury

The above observation led us to examine whether autophagy impairment due to DDC exposure can exacerbate p62-IHB accumulation to aggravate cholestatic liver injury.DDC diet was fed to Atg7+/-mice for 2 weeks. Atg7 is an important autophagy-related gene required for the formation of autophagosomes[2]. We choose hepatocyte specific Atg7+/-in place of Atg7-/-mice for DDC treatment because Atg7-/-develop cholestatic liver injury on its own[37-39]. The gross examination revealed enlarged and dark brown livers DDC exposed Atg7+/-similar to its wild type counterpart (**Fig.6A**).Interestingly, DDC-mediated hepatomegaly was further exacerbated in the Atg7+/- liver(**Fig.6B**). More importantly, the DDC exacerbated the cholestatic liver injury as examined by serum ALT, ALP, TBA, TB, DB in Atg7+/- mice(**Fig.6C, Supplementary Fig.6A**).General histological and immunohistochemistry examination also presented increased immune cells infiltration, and ductular reaction in Atg7+/- mice when compared to DDC fed wild type mice(**Fig.6D, Supplementary Fig.6B**). Moreover, examination of p62 and IHB related proteins showed furthermore accumulation of IHBs related proteins in DDC fed Atg7+/- mice(**Fig.6E**).Immunofluorescence staining for K8/K18 protein showed linear distinct, bright filamentous staining pattern that runs along the cell periphery in Atg7+/- mice(**Fig.6F**). This K8/K18 immunostaining pattern is distinctly different when compared to diffused staining pattern of DDC excposed wild type liver(**Fig.6F**).Thus genetic impairment of autophagy function exacerbates DDC induced cholestatic liver injury.

**Figure 6.**
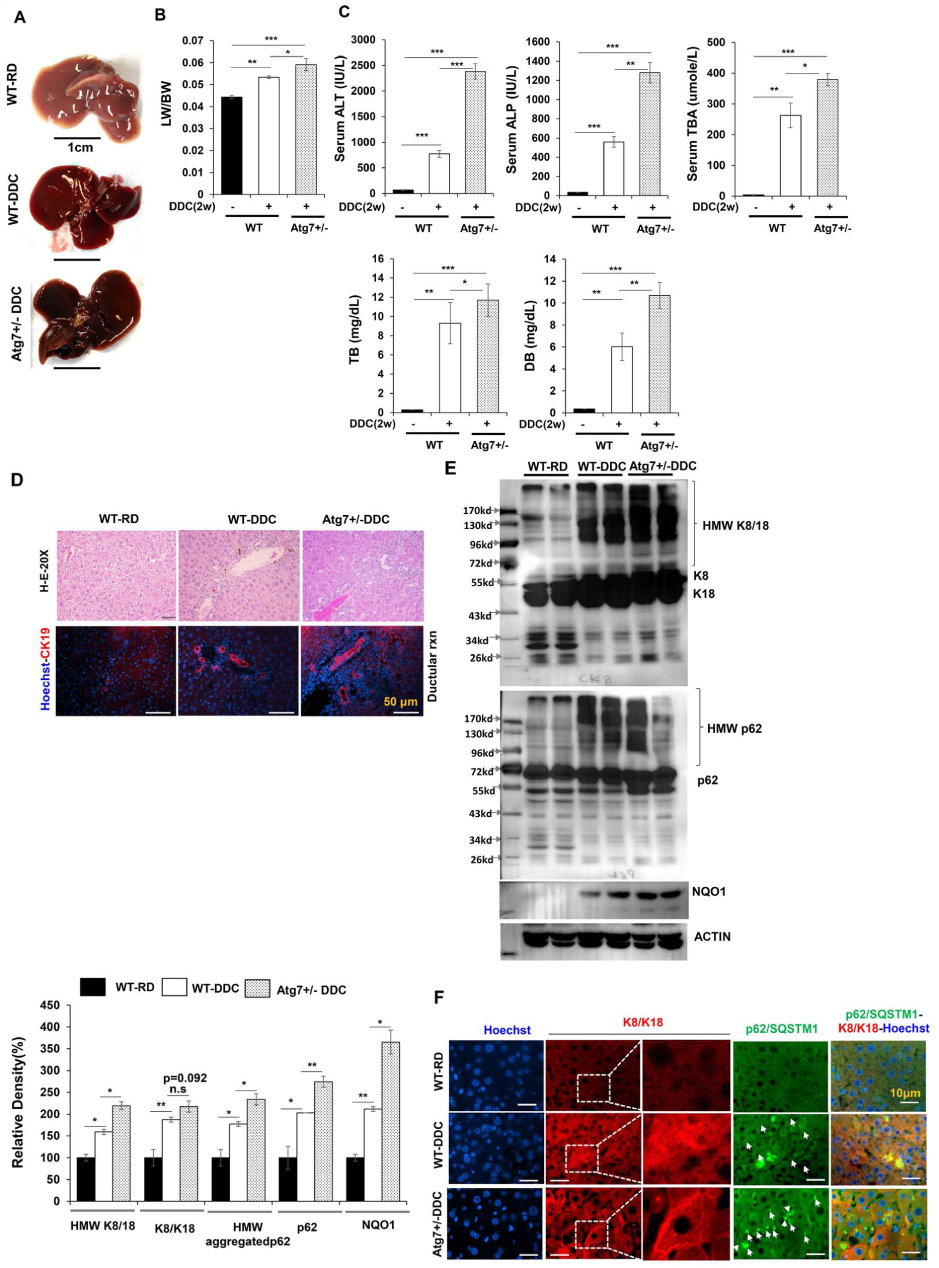
Impaired hepatic autophagy exacerbates DDC mediated cholestatic liver injury. (A) Representative gross morphology of liver of regular diet(RD) or DDC diet fed wild type mice and 2-week DDC diet fed Atg7+/-mice. (B) LW/BW ratio showing hepatomegaly in the 2 weeks DDC diet fed wild type or Atg7+/-mice. (C) Serum ALT, ALP, total Bile Acid(TBA), Total bilirubin(TB), and direct bilirubin(DB) levels were quantified in the 2 weeks DDC diet fed wild type or Atg7+/-mice. Data are expressed as the mean ± SEM. n.s: not significant,**P≥0.01, ***P≥0.0001 (n=3). (D) Liver sections were subjected to H&E staining (original magnification, ×200), and immunostained for CK19 to detect Ductular cells(original magnification, ×200). Scale bars: 50 μm (CK19).(E) Total liver lysates were analyzed by immunoblotting for K8/18, p62, and GAPDH proteins in the 2 weeks DDC diet fed wild type or Atg7+/-mice. Protein band intensity was normalized to GAPDH band intensity. Data are expressed as the mean ± SEM. n.s: not significant,*P≥0.05, ***P≥0.001 (n=2). (F) Liver sections were co-stained for K8/K18 and p62. Scale bars: 10 μm.

### 7. DDC-mediated autophagy impairment activates p62-NRF2 and mTORC1 signaling but not the NF-kB pathway

Hepatic autophagy impairment due to DDC exposure cause accumulation of p62(**Fig.1D,1E, 1F, 1H**). It is well established that hepatic p62 not only acts as an autophagy substrate but can functions as a signaling hub[40]. The p62 is a stress-inducible and multifunctional protein that contains several protein-protein interaction domains-a Phox1 and Bem1p (PB1) domain, a zinc finger (ZZ), two nuclear localization signals (NLSs), a TRAF6 binding (TB) domain, a nuclear export signal (NES), and LC3-interacting region (LIR), a Keap1-interacting region (KIR), and a ubiquitin-associated (UBA) domain[41, 42]. These p62’s protein domains can interact with various binding partners to activate downstream signaling pathways and may play a pathogenic role in DDC exposed liver. We next examined the potential signaling pathways specifically nuclear factor kappa B(NF-kB), mammalian target of rapamycin complex 1 (mTORC1), and NRF2 signaling pathways that could have pathological impact in DDC exposed liver.

P62 regulates NF-kB by interacting with aPKC through its PB1 domain, RIP through its ZZ domain, and TRAF6 through its TB domain [43]. As such, the p62 protein interacts with RIP and bridges RIP to aPKCs, resulting in the activation of NF-kB by the TNFa signaling pathway [44]. Interestingly, the expression level of TNFa has been reported to be elevated in DDC conditions [9]. Hence, NF-kB pathway could be activated, as a minor inflammatory response in DDC exposed liver. So, we examine whether the accumulation of p62 in DDC causes activation of the NF-kB inflammatory signaling pathway.Immunoblot analysis showed a lower level of p65/NF-kB protein in DDC diet-fed mice (**Fig.7A**). the mRNA expression analysis for NF-kB downstream target genes such as c-Rel, Cox2, Rel B, or NF-kBIZ were also significantly downregulation(**Fig. 7B**) suggesting that the NF-kB pathway is not activated rather suppressed with an acute DDC exposure.

**Figure 7.**
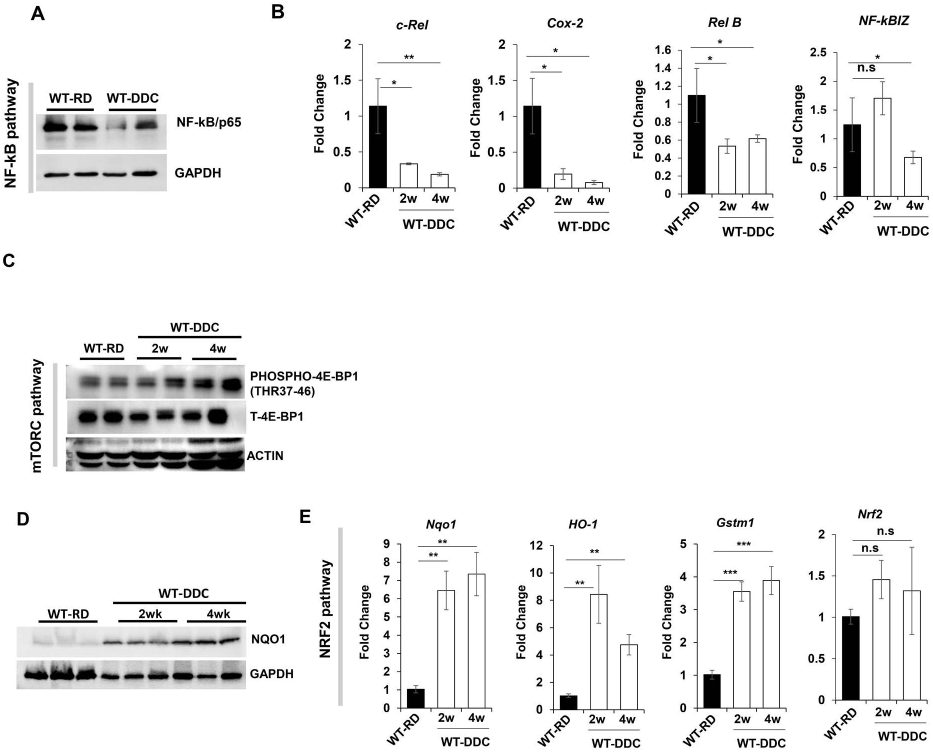
Autophagy impairment is linked to activation of the NRF2 and mTORC1 signaling pathways, not to the NF-kB signaling pathway. (A) Total liver lysates were analyzed by immunoblotting for p65/NF-kB and GAPDH proteins in wild type mice fed with regular diet(RD), and 2 weeks of DDC diet. (B) Quantitative PCR analysis for c-Rel, COX2, Rel-B and NF-kBIZ mRNA expression in wild type mice fed with RD, and 2-4 weeks of DDC diet. The mRNA expression levels were normalized to actin. Data are expressed as the mean ± SEM. n.s: not significant, *P≥0.05, **P≥0.01,n=3). (C-D) Total liver lysates were analyzed by immunoblotting for (C) phospho-4E-BP1, Total 4E-BP1, and ACTIN proteins and (D) NRF2, NQO1, and GAPDH in wild type mice fed with RD, and 2-4 weeks of DDC diet. (E) Quantitative PCR analysis for Nqo1, HO-1, Gstm1, and Nrf2 mRNA expression in wild type mice fed with RD, and 2-4 weeks of DDC diet. Data are expressed as the mean ± SEM. n.s: not significant,**P≥0.01, ***P≥0.0001 (n=3).

P62 can also promote mTORC1 activation by directly interacting with Raptor, a key component of mTORC1 [45]. The region between the ZZ and TB domains (amino acids 167–230) of p62 is required for the interaction between p62 and Raptor [45]. Moreover, over-expression of p62 enhanced mTORC1 activation [45]. Immunoblot analysis of mTORC1 downstream target protein showed elevated levels of phosphorylated-4E-BP(**Fig. 7C**) suggesting that p62 accumulation can activate the mTORC1 pathway in the DDC diet. The activation of the mTORC1 pathway could further suppress autophagy causing a vicious cycle of autophagic suppression.

Accumulation of p62 can also activate NRF2 by a non-canonical pathway [46, 47]. The p62 protein competitively binds to Kelch-like-ECH-associated protein 1 (KEAP1), an adaptor component for Cullin3-based E3 ubiquitin ligase complex that ubiquitinates NRF2 for proteasomal degradation. Moreover, NRF2 is an anti-oxidative stress-activated transcription bZIP factor, that can promote the transcriptional expression of K8/K18 expression[13].On the other hand, DDC-mediated inhibition of ferrochelatase causes accumulation of PP-IV that can cause oxidative stress [48] and hence could activate NRF2. So we next analyzed the expression level of NRF’s downstream target Protein NQO1 in DDC exposed liver.Immunoblot analysis showed marked elevation of NQO1 in DDC exposed liver (**Fig. 7D**). Activation of NRF2 transcription factor cause nucleus translocation where it binds to the anti-oxidative response element present in the promoter region of a battery of anti-oxidative response genes such as NQO1, HO-1, and Gstm1. Quantitative PCR analysis showed that NRF2 downstream target genes NQO1, HO-1, and Gstm1 also significantly upregulated in DDC exposed liver. (**Fig. 7E**). Notably, the NRF2 mRNA expression did not change, suggesting that NRF2 protein accumulation and its activation is via non-canonical p62 accumulation as observed in the DDC diet exposed liver. Overall these results clearly suggest that accumulation of p62 due to autophagy impairment in DDC diet leads to activation of NRF2 and mTORC1 signaling pathway. The NF-kB signaling pathway is not activated, but rather suppressed in the liver treated with a DDC.

### 8. Impaired autophagy by DDC is associated with suppression of hepatic FXR signaling and deregulated bile acid (BA) metabolism

Our previous study shows that autophagy-deficiency in hepatocyte deregulated BA metabolism resulting in an intracellular cholestasis with increased levels of serum and hepatic BA level[37]. Autophagy-deficiency causes the accumulation of p62, activating the non-canonical NRF2 pathway which then transcriptionally represses the expression of FXR nuclear receptor, a master regulator of hepatic BA metabolism [37]. It is well established that DDC exposed liver displays BA metabolic disturbance and intrahepatic cholestasis by an unclear mechanism [9]. Since DDC impairs hepatic autophagy and activates the NRF2 signaling, we next examined whether activated NRF2 in DDC could also relate to FXR downregulation together with other abnormalities related to BA metabolism.

There was a marked decrease in the FXR protein level in the DDC exposed liver compared to wild type control liver(**Fig.8A**). The mRNA level of FXR and its downstream target genes-SHP, BSEP, Osta were also significantly downregulated(**Fig.8B**). However. mRNA expression level of RXRa, which is a binding partner of FXR did not show significant changes in the DDC exposed liver(**Fig.8B**). Additionally, serum levels of Total Bile acid(TBA), cholesterol, triglyceride, Total Bilirubin, Direct bilirubin, and liver injury markers-ALT and ALP were dramatically upregulated reflecting the ongoing cholestatic liver injury in the DDC exposed mice (**Fig.8C, Supplementary Fig.7A-7B**). Interestingly the cholestatic liver injury parameters were partially recovered with prolonged DDC exposure suggesting the development of a compensatory resistance mechanism in the liver(**Fig.8C, Supplementary Fig.7A-7B**).

**Figure 8.**
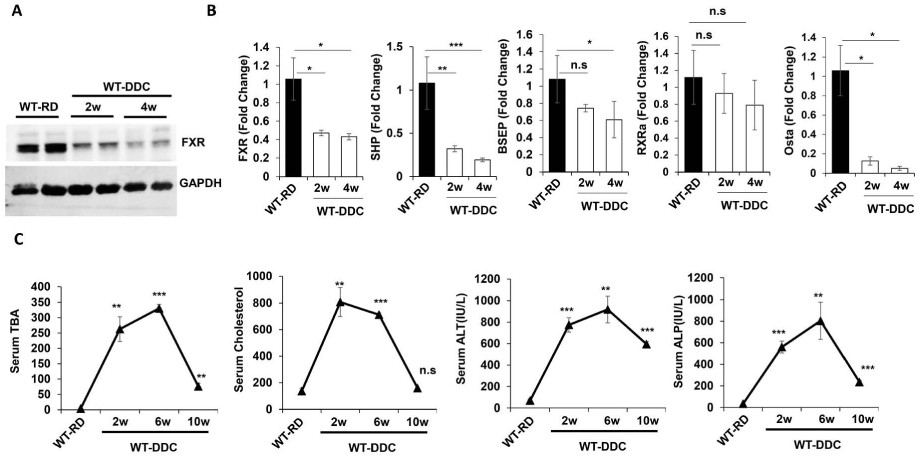
DDC repress hepatic FXR signaling and impairs bile acid handling. (A) Total liver lysates were analyzed by immunoblotting for FXR and GAPDH proteins in wild type mice fed with regular diet(RD), and 2-4 weeks of DDC diet. (B) Quantitative PCR analysis for FXR, SHP, BSEP, RXRa, and OSTa mRNA expression in wild type mice fed with RD, and 2-4 weeks of DDC diet. The mRNA expression levels were normalized to actin. Data are expressed as the mean ± SEM. n.s: not significant, *P≥0.05, **P≥0.01, ***P≥0.0001,(n=3). (C) Serum total Bile Acid(TBA), Cholesterol. ALT, and ALP levels were quantified for 9-week-old mice fed with RD, and 2-10 weeks of DDC diet.Data are expressed as the mean ± SEM. n.s: not significant,**P≥0.01, ***P≥0.0001 (n=3).

Expression analysis of various BA transporters showed elevation of Apical (MDR1A, MDR1B) and systemic (Ostb, Mrp3, Mrp4) BA transporters.The expression levels of basolateral or enterohepatic (NTCP, OATP2, OATP1, and OATP4) BA transporters were suppressed(**Supplementary Fig.8**) indicating the compensatory adaptation in responses to cholestasis. Taken together, these results suggest that DDC causes downregulation of FXR nuclear receptors and cholestatic liver injury. DDC diet-activated p62-Nrf2 signaling pathway may suppress FXR and cause cholestatic liver injury.

## DISCUSSION

### A. Role of Autophagy in acute DDC associated IHBs formation

Hepatocyte protein aggregates, including MDBs, are usually seen in hepatocellular carcinoma, Wilson’s disease, non-alcoholic steatohepatitis, and several other chronic liver disorders. MDBs are primarily composed of K8/18 as well as p62, Ub, and chaperones[12, 36]. IHBs contain p62 and ubiquitin (although not constantly) but differ from MDBs by lacking K8/K18[5]. Incorporating “abnormal” keratins and HSPs into aggregated p62 causes p62-containing IHBs to become classical MDBs [5, 22]. MDBs can be experimentally induced in the livers of mice chronically fed DDC[5]. The cellular status of these established MDB proteins is less clear in the acute model of DDC intoxication. We show that an acute DDC exposed liver has an increased Ub and p62 positive protein aggregates, as seen in autophagy-deficient livers(**Fig.1D-1F, 1H**)[39]. The p62 aggregate formation could be caused by impaired autophagy clearance. Autophagic degradation is involved in the disposal of aggregated proteins. The DDC diet suppressed autophagic activity and thus compromised the clearance of IHBs. As a result of inadequate protein degradation via autophagy, IHBs accumulate.

Various HSPs that are involved in regulating protein conformation is significantly decreased in the liver of DDC intoxicated mice (**Fig.2A-2J**). Thus, despite elevated K8/K18 protein levels and p62-Ub-containing IHBs, MDBs were not formed in acute DDC intoxicated livers. The lack of MDB formation at the early stages of DDC intoxication could be due to HSPs downregulation. What causes the downregulation of so many HSPs in the liver of DDC diet-exposed mice is unclear.However, in situ promoter analysis with MatInspector and JASPER revealed the presence of putative anti-oxidative response element(ARE) binding site to NRF2 in HSF1, the master regulator of the expression of HSPs (Katherine B et al unpublished data). One possible explanation would be the activation of NRF2 could suppress HSF1 and therefore downregulate HSPs (**Fig.9**)

**Figure 9.**
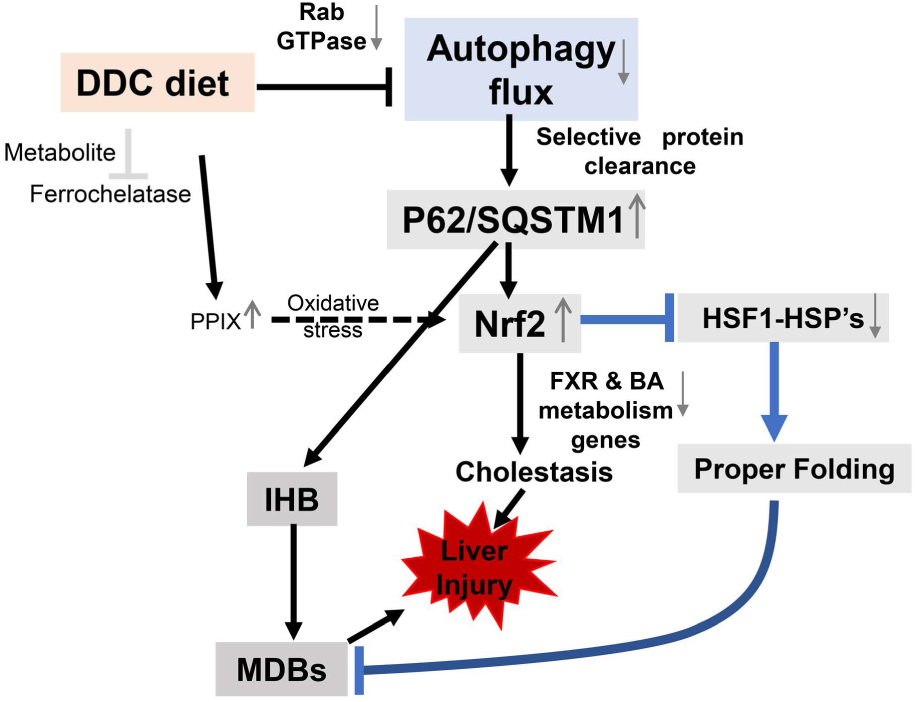
Schematic model showing role of autophagy in DDC diet. Acute DDC impairs the autophagic flux due to Rab suppression. Impaired autophagy leads to formation of p62 containing intrahyaline bodies(IHB). Impaired autophagy can also relate to downregulation of hepatic protein chaperonin system, further aiding in IHB formation. Further activation of P62 related NRF2 can suppress the FXR nuclear receptor and cause cholestatic liver injury.

### B. Role of Rabs in autophagy flux impairment

The Rab proteins are members of the Ras superfamily of small G proteins. They function as regulators of vesicular traffic by switching between GTP-bound and GDP-bound conformations. Rab bound to GDP is inactive, whereas Rab bound to GTP is active. GDP/GTP exchange factors (GEPs), GDP dissociation inhibitors (GDIs), and GTPase-activating proteins (GAPs) regulate the conversion from one state to another. Rab proteins are found in distinct subcellular locations. The Rab7 is present in autophagosomes and [21, 36] promotes microtubule plus-end-directed transport and fusion of autophagosomes with lysosomes through a novel FYVE and coiled-coil domain-containing protein FYCO1[20]. Rab7 is also involved in the maturation of late autophagic vacuoles, recruitment of dynein/dynactin motors, Rabring7 (Rab7-interacting ring finger protein), and the hVPS34/p150 complex[36]. Decreased expression of Rab proteins could mechanistically explain why DDC impairs autophagosomes and lysosomal fusion. Since Rab proteins are also localized to the conventional secretory pathway, DDC may compromise the hepatic conventional secretory pathway.

It is unclear why the DDC diet causes massive repression of Rab proteins. There is a possibility that DDC suppresses upstream regulators of Rab expression. Alternatively, DDC-related metabolites generated in the liver may inhibit Rab expression. DDC N-alkylates the heme of certain hepatic cytochrome-p450 enzymes to produce N-methyl protoporphyrin(NMPP). NMPP is a potent inhibitor of ferrochelatase which is involved in the conversion of PP-IX into heme, thus resulting in the accumulation of PP-IX and upstream porphyrin intermediates[37]. The accumulation of NMPP or PP-IX could be responsible for the downregulation of Rab proteins. A future study will examine the common motifs present in the promoters of the Rab genes and the transcription factor that regulates their expression.

### C. Autophagy regulates hepatic xenobiotic metabolism by modulating the p62-NRF2-FXR signaling

A xenobiotic like DDC may interfere with cell survival mechanisms such as autophagy. Our results indicate that the acute DDC exposure impairs fusion of autophagosome with lysosome causing p62 accumulation and IHB formation. It is unclear how exactly this fusion of autophagosome and lysosome is impaired by DDC. However, this study support our previous study showing that autophagy regulates signaling proteins such as p62 and NRF2[35, 36]. Autophagy is necessary for maintaining FXR functionality, and a deficiency of autophagy suppressed its expression via NRF2 activation. The accumulation of p62 in the autophagy-deficient liver causes NRF2 activation and FXR repression, resulting in cholestatic liver injury[37]. We confirm the pathological significance of the p62-NRF2-FXR signaling axis in the DDC model of xenobiotic-induced liver injury. As an alternative, DDC can cause oxidative stress, resulting in the oxidation of cysteine residues in KEAP1, which then activates NRF2 via conventional pathway[49]. DDC feeding may also inhibit NRF2 proteasomal degradation, most likely through PpIX-mediated oxidative stress[37, 38].

Moreover, the role of autophagy in xenobiotic-induced liver injury is unclear. Our data strongly suggest that mice with impaired autophagy such as Atg7+/-mice are more susceptible to DDC-induced liver injury than those fed the regular diet. The DDC diet injury is cholestatic since there is an elevation of serum total bile acid in DDC diet animals. This implies that autophagy function impairment could exacerbate the DDC diet injury, and activating autophagy may represent an effective therapeutic approach.

## SUPPLEMENTARY FIGURES

**Supplementary Figure 1.**
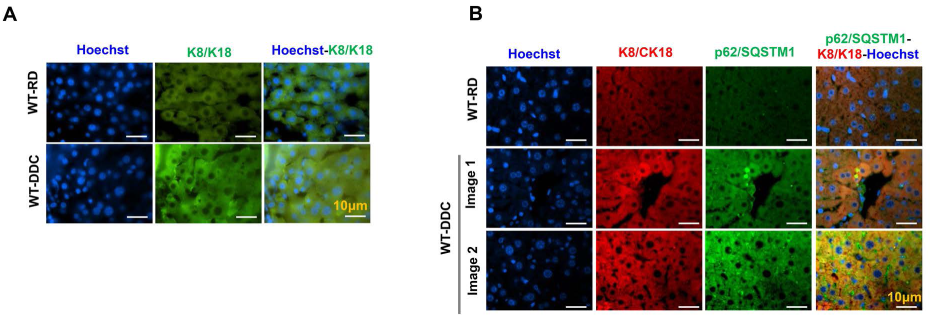
Hepatic IHB formation in DDC diet exposed liver. (A) Liver sections were stained for K8 and K18. (B) Co-immunostaining for K8/K18 and p62 in liver section from liver section of regular diet or DDC diet fed wild type liver. Scale bars: 10 μm

**Supplementary Figure 2.**
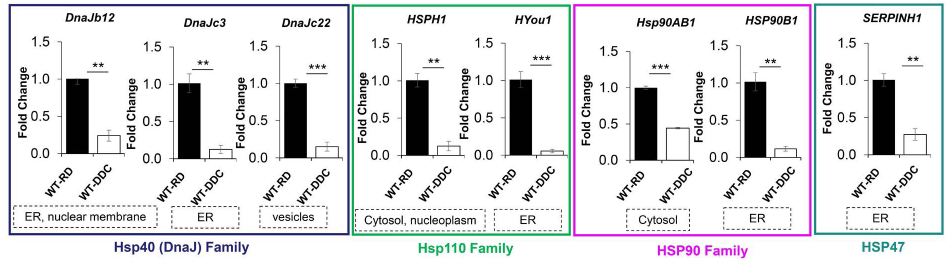
HSPs are downregulated regardless of subcellular localization. (A) Quantitative PCR analysis for mRNA expression of HSP’s localized to endoplasmic reticulum(ER), nuclear membrane, vesicles, cytoplasm, nucleoplasm, ER that belongs to Hsp40 (DnaJ) Family, Hsp110 Family, HSP90 Family, or HSP47 Family. The cDNA was prepared from mRNA isolated from 9-week-old mice wild type mice fed with regular diet(RD) and 2 weeks of DDC diet.The mRNA expression levels were normalized to actin. Data are expressed as the mean ± SEM. n.s: not significant,*P≥0.05, **P≥0.01, ***P≥0.001 (n=3)

**Supplementary Figure 3.**
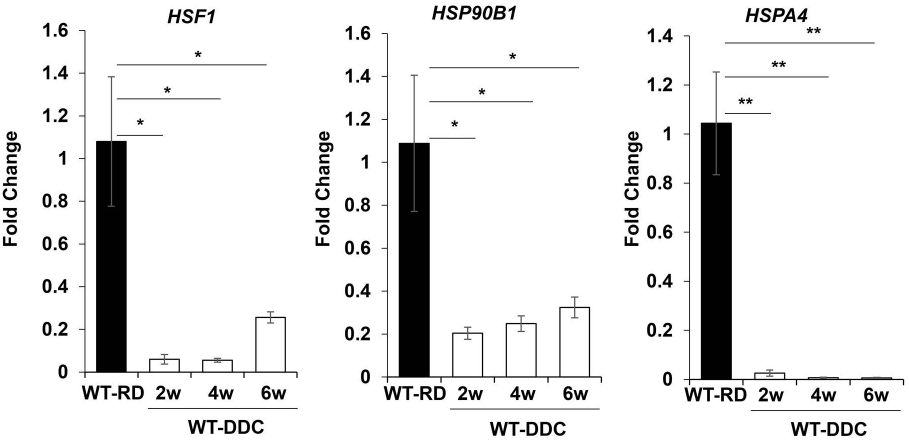
HSF1 and HSP’s remain suppressed in longer DDC fed liver. Quantitative PCR analysis for HSF1,HSP90B1, and HSPA4 mRNA expression from 9-week-old mice wild type mice fed with regular diet(RD) or 2-6 weeks of DDC diet.The mRNA expression levels were normalized to actin. Data are expressed as the mean ± SEM. *P≥0.05, **P≥0.01,(n=3).

**Supplementary Figure 4.**
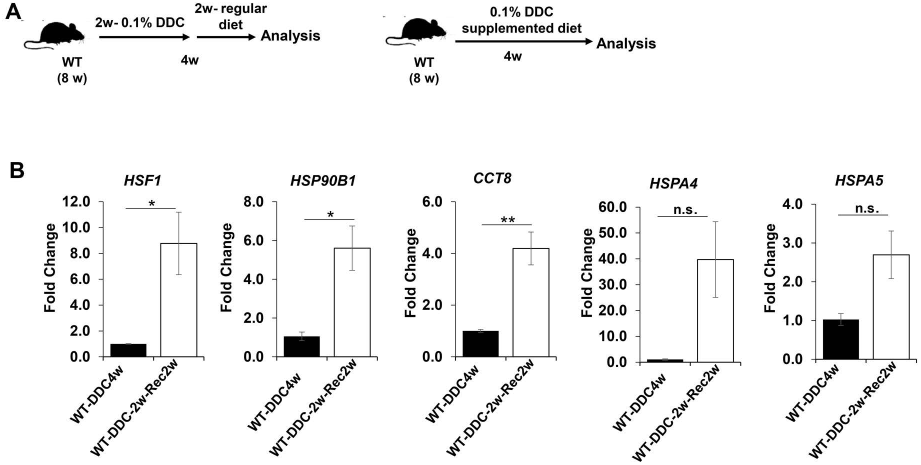
Hepatic suppression of HSF1 and HSP is reversible. (A) Schematics of DDC diet recovery experimental model.(B) Quantitative PCR analysis for HSF1, HSP90B1,CCT8,HSPA4 and HSPA5 mRNA expression from 8-week-old mice wild type mice fed with either 4 weeks of DDC or 2 weeks DDC diet followed by 2 weeks Regular Diet(RD) The mRNA expression levels were normalized to actin. Data are expressed as the mean ± SEM. n.s: not significant, *P≥0.05, **P≥0.01,(n=3).

**Supplementary Figure 5.**
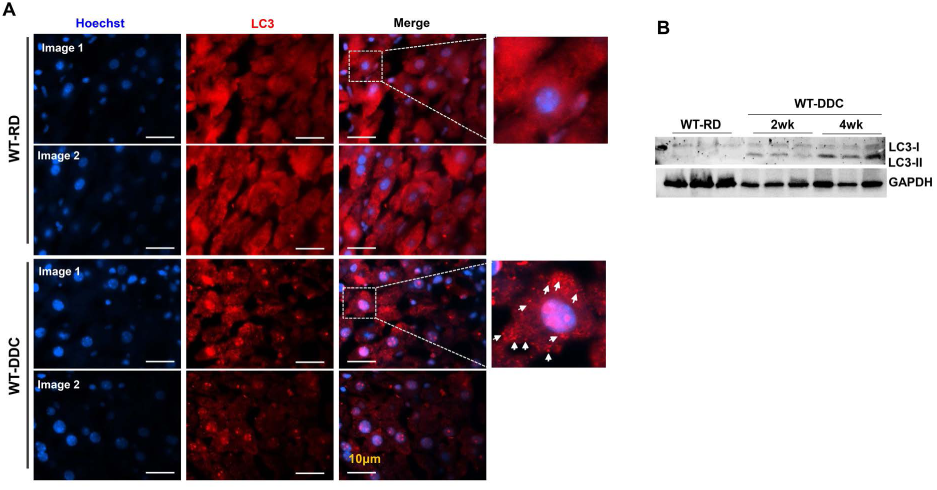
Autophagy markers are elevated in DDC diet exposed liver. (A) Autophagosomes detection by LC3 Immunostaining in liver section prepared from wild type mice fed with regular diet(RD) and 2 weeks of DDC diet. Scale bars: 10 μm. (B) Total liver lysates prepared from RD or 2-4 weeks of DDC diet fed wild type mice, were analyzed by immunoblotting for LC3, and GAPDH.

**Supplementary Figure 6.**
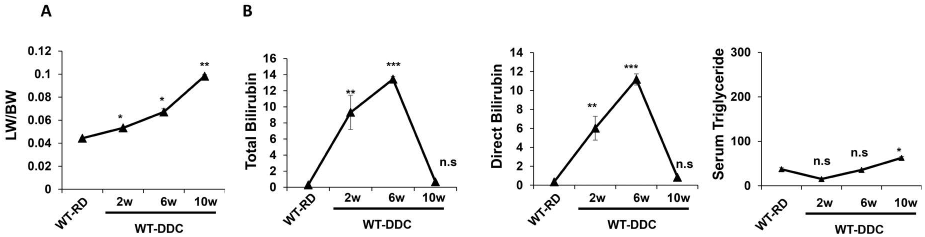
Cholestasis related parameters analysis in DDC diet exposed mice. (A) Serum total cholesterol(TC), and triglyceride levels were quantified in the 2 weeks DDC diet fed wild type or Atg7+/-mice. Data are expressed as the mean ± SEM. n.s: not significant,**P≥0.01, ***P≥0.0001 (n=3). (B) Liver sections were subjected to H&E staining (original magnification, 100X and ×200).

**Supplementary Figure 7.**
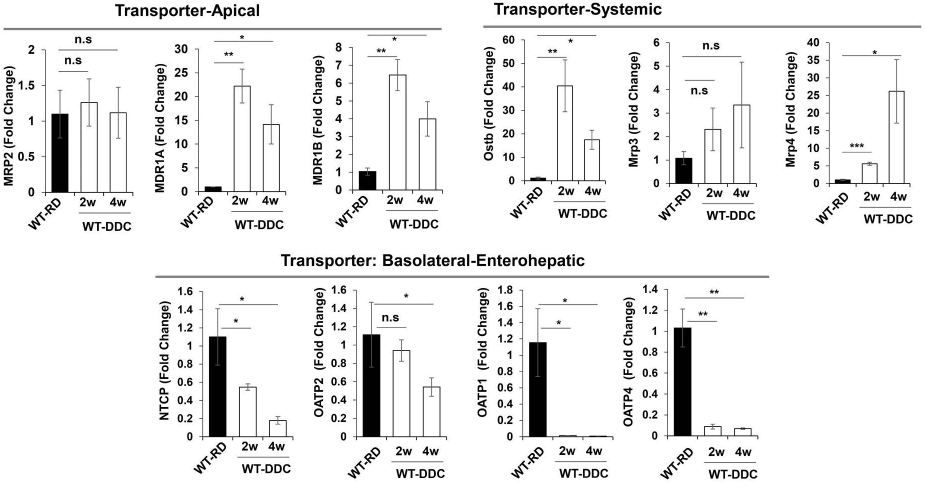
DDC cause hepatomegaly and cholestatic liver injury. (A) LW/BW ratio showing hepatomegaly in the DDC diet fed wild type mice for different time period.(B) Serum total bilirubin, Direct bilirubin and triglyceride levels were quantified for 9-week-old mice fed with regular diet(RD), and 2-10 weeks of DDC diet. Data are expressed as the mean ± SEM. n.s: not significant,**P≥0.01, ***P≥0.0001 (n=3).

**Supplementary Figure 8.**
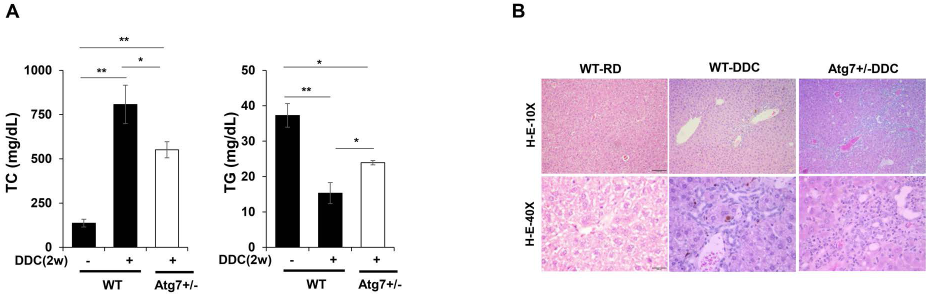
Adaptation alteration in cholestasis associated bile transporters in DDC diet. Quantitative PCR analysis for Apical Bile acid transporters(MRP2, MDR1A, MDR1B), Systemic BA transporters(OSTb, MRP3, and MRP4), and Basolateral/Enterohepatic BA transporters(NTCP, OATP2, OATP1, and OATP4) mRNA expression in wild type mice fed with regular diet(RD), and 2-4 weeks of DDC diet. The mRNA expression levels were normalized to actin. Data are expressed as the mean ± SEM. n.s: not significant, *P≥0.05, **P≥0.01, ***P≥0.0001,(n=3).

## Acknowledgments

We acknowledge the support of This study was supported in part by the Lousiana Board of Regent(BK),BeHEARD Rare disease(BK), ASIP/SROPP(BK),and Tulane School of Medicine startup fund(BK) and NIH/NIDDKD grants R01 DK116605 (XMY).We also thank the support of the Tulane University Histology section at the TUSOM Department of Pathology and Laboratory Medicine for liver tissue sample processing, general histological staining, and providing unstained slides. We thank all members of Dr. Khambu’s and Dr.Yin’s laboratory for critical discussion about the xenobiotic metabolism and hepatic autophagy.

