## Supplementary Table 1-2 for "Impaired hepatic autophagy exacerbates xenobiotics induced liver injury"

**Supplementary Table 1**: RT-qPCR primers

### Supplementary Table 1. Primers used in PCR assays

| Gene Name | Forward Primer | Reverse Primer |
|---|---|---|
| BSEP | 5'-CCTCTCACCAGGCTCTCTACC-3' | 5'-CGCCACTGTGGAAAGTCAGGG-3' |
| CCT2 | 5'-TCACCACAAGGACCACTTTAC-3' | 5'-CAGACTCCCACCTAGTTTCTTG-3' |
| CCT3 | 5'-GACCTGCTTGGGACCTAAAT-3' | 5'-GGATGCTGGACTTGAATCTCT-3' |
| CCT4 | 5'-TCCTACTGTGTTCGTGCTTTC-3' | 5'-GGCGGTTTCTTAGCTCTGTTA-3' |
| CCT5 | 5'-GCTGGGCTCCAAAGTGATTA-3' | 5'-CAACATCTCTCCGCTCCATATC-3' |
| CCT6A | 5'-GTCTACCCTTGTTCCGTAAGTG-3' | 5'-CTGTAGCCTGAGTTGGCATAG-3' |
| CCT7 | 5'-CCGAGGCAAAGCAACAATATC-3' | 5'-GGACTTGGCTATGTCCACTAAA-3' |
| CCT8 | 5'-CTGTGTACTCTTGTCCGTTTGA-3' | 5'-TGAGGTTCTCCTCTCCCTTAC-3' |
| CHOP | 5'-TCTGATTGACCGAATGGTGA-3' | 5'-TCTGGGAAAGGTGGGTAGTG-3' |
| K18 | 5'-GGACATCGAGATCACCACCT-3' | 5'-TGAAGCCAGGGCTAGTGAGT-3' |
| K8 | 5'-CAAGTCTGCCGAAATCAGGGAC-3' | 5'-TCCAAGTTGATGTTCTGGTTTT-3' |
| COX2 | 5'-TGCAGAATTGAAAGCCCTCT-3' | 5'-CCCCAAAGATAGCATCTGGA-3' |
| c-Rel | 5'-AGTGACTCACCCCACCTCAC-3' | 5'-AGGCCCTTCTAGGAATGGAA-3' |
| DnaJa3 | 5'-CATCCACTCGGACCTCTTTATTT -3' | 5'-GCAGGGATCGTCACATTGAT-3' |
| DnaJb12 | 5'-GTCTCCAGGCTTATGGTTAAGG-3' | 5'-TGTCTGGGATCTCTGCTGTA-3' |
| DnaJc22 | 5'-CTTGTCAGCACCATCCTCAA-3' | 5'-CTCTACCCTTCTCATCCCTGTA-3' |
| DnaJc3 | 5'-AGCAAGGAGCTAACCGTATTT-3' | 5'-CTGAACACCTCCACAGGATAAG-3' |
| FXR | 5'-GGCCTCTGGGTACCACTACA-3' | 5'-TGTACACGGCGTTCTTGGTA-3' |
| Gstm1 | 5'-ACTTGATTGATGGGGCTCAC-3' | 5'-TCTCCAAAATGTCCACACGA-3' |
| HO-1 | 5'-GTGATGGAGCGTCCACAGC-3' | 5'-TTGGTGGCCTCCTTCAAGG-3' |
| HSF1 | 5'-GTTCCAGCATCCTTGTTTCTTG-3' | 5'-GACACTGTCCTGGCGTATTT-3' |
| HSP47= SERPINH1 | 5'-CATCTTCCTGGTGCGAGATAA-3' | 5'-CCACTCTTGGACTCTACAACTC-3' |
| Hsp90AB1 = HspC3 | 5'-GCTATCCCATCACCCTCTATTT-3' | 5'-GCTTCTCCTCATCCTCCTTATC-3' |
| Hsp90b1 = HspC4 | 5'-GCCCTCAAGGACAAGATAGAAA-3' | 5'-TGTTGCCAGACCATCCATAC-3' |
| HSPA1A | 5'-TGGTTGCACTGTAGGACTTG-3' | 5'-CGAGTTCAGGATGGTTGTGT-3' |
| HSPA4 | 5'-TCTGAGCAGTCCATCCTTAGTA-3' | 5'-GAACTCTCATCCTGTCCCATTC-3' |
| HSPA5 | 5'-GAGACTGCTGAGGCGTATTT-3' | 5'-TGACATTCAGTCCAGCAATAGT-3' |
| HSPA8 | 5'-ACTCCTCTTTCCCTTGGTATTG-3' | 5'-GTCAGAGTAGGTGGTGAAAGTC-3' |
| HSPA9 | 5'-ACTCCTGTGTGGCTGTTATG-3' | 5'-GTCGTTCTCCATCTGCTGTAA-3' |
| HSPD1 | 5'-AGGTTGTGAGAACTGCCTTAC-3' | 5'-TCCAGGGTCCTTCTCTTCTT-3' |
| HSPE1 | 5'-ACTGTAACCAAAGGTGGCATTA-3' | 5'-GGCTCAATCTCTCCACTCTTTC-3' |

| | | |
|---|---|---|
| Hsph1 | 5'-TCTTCAGTGTGGAGCAGATAAC-3' | 5'-GAAGAAGGATGGGACTGAGATG-3' |
| HYou1 | 5'-CCCAGAATCTGACCACAGTAAA-3' | 5'-GCCACTCTCATCCAGGTTAAA-3' |
| MDR1A | 5'-AAAGGCTCTACGACCCCCTA-3' | 5'-CCTGACTCACCACACCAATG-3' |
| MDR1B | 5'-TTGGTGGCACAACAACTCAT-3' | 5'-GGCTTTCGCATAGTCAGGAG-3' |
| MRP2 | 5'-GCACTGTAGGCTCTGGGAAG-3' | 5'-TGCTGAGGGACGTAGGCTAT-3' |
| Mrp3 | 5'-GGACTTCCAGTGCTCAGAGG-3' | 5'-AGCTGTGGCCTCGTCTAAAA-3' |
| Mrp4 | 5'-TGTTTGATGCACACCAGGAT-3' | 5'-GACAAACATGGCACAGATGG-3' |
| Nf-KBIZ | 5'-GTGGAGGCAAAGGATCGTAA-3' | 5'-TCACGAAAGACAGGCAACTG-3' |
| Nqo1 | 5'-AGCGTTCGGTATTACGATCC-3' | 5'-AGTACAATCAGGGCTCTTCTCG-3' |
| NTCP | 5'-CACCATGGAGTTCAGCAAGA-3' | 5'-CCAGAAGGAAAGCACTGAGG-3' |
| OATP1 | 5'-ATCCAGTGTGTGGGGACAAT-3' | 5'-GCAGCTGCAATTTTGAAACA-3' |
| OATP2 | 5'-TTGCTGACTGCAACACAAAG-3' | 5'-TGGTTCCAGTTCCAACAGAC-3' |
| OATP4 | 5'-TGGGATTCCATTCACTGGTT-3' | 5'-TGCTCCACAGCTGGTTACAG-3' |
| Osta | 5'-GTCTCAAGTGATGAACTGCCA-3' | 5'-TTGAGTGCTGAGTCCAGGTC-3' |
| Ostb | 5'-GTATTTTCGTGCAGAAGATGCG-3' | 5'-TTTCTGTTTGCCAGGATGCTC-3' |
| p62/SQSTM1 | 5'-GCTCAGGAGGAGACGATGAC-3' | 5'-AGAAACCCATGGACAGCATC-3' |
| Rab 13 | 5'GAGATCGGGAACCAACAGTAAG -3' | 5'-GGGTGAATAGGAGGCAAGAAA-3' |
| Rab10 | 5'-AACCTCCTAACCTGGATTTGAC-3' | 5'-CCCACTCCTTCTTGCTCTTT -3' |
| Rab11b | 5'-CCCAGCTCTCGAACTCTTATTC-3' | 5'-CAGAAGCTGAGTGGTAGGTTTC-3' |
| Rab12 | 5'-AAGGAGACGTTCGATGACTTG-3' | 5'-CTGTCTCACAGTCCAGCTTATT-3' |
| Rab14 | 5'-CACACACGGAAATAAGACACAAC-3' | 5'-CAGGAGCTCTTTCCAGCATTA-3' |
| Rab17 | 5'-CTGTAATCACTGCTTGCCAAAG-3' | 5'-TCTGGGTATCAGGTAAGGTAGG-3' |
| Rab18 | 5'-GCATCCCAGAACTCACCTAAA-3' | 5'-GCTTGATTCTGGAGCCTCTATC-3' |
| Rab1b | 5'-CATGGCATCATTGTGGTGTATG-3' | 5'-TGTTGCCTACCAGGAGTTTATT-3' |
| Rab20 | 5'-GGCCGCTATCATCCTTACATAC -3' | 5'-CAGTCATTGTTGGCTGTTTCTG-3' |
| Rab21 | 5'-GGAGCCAAGCATTACCATACT-3' | 5'-CTGGGCTGTCTCTATCATTCTTT-3' |
| Rab22a | 5'-CAGCTTCCGCCATCTCTAAA-3' | 5-CGTGGGTTCTGACACATACA-3' |
| Rab23 | 5'-CCGACAGGTAGATGAAAGGAATG-3' | 5'-GCTTACAGTGGCTATGGAGAAG-3' |
| Rab28 | 5'-GCTCTGCTCCTTCATCCTTTAG-3' | 5'-ATGGACTCTCCATTCGGATTTC-3' |
| Rab29 | 5'-CACATCCATGACACGACTCTAC -3' | 5'-GTCCAGATCCTGTTTCCATCTT-3' |
| Rab2a | 5'-GCAGGAGTCCTTTCGTTCTATC-3' | 5'-GTTGAACGTGTCTCTCCTTGT-3' |
| Rab31 (Rab22b) | 5'-CTCAAGACCATCAGTGCCTATC-3' | 5'-TCAGCTCAATCGTTCTCTGTTT-3' |
| Rab34 | 5'-CTCCTGCTCAGTATTCCCTAATG-3' | 5'-CTGCCACACGGAAGAAGAA-3' |
| Rab35 | 5'-CCACATCGGGCTCAGTATTT-3' | 5'-ACAGGTATGAGGGTGCAAAG-3' |
| Rab3a | 5'-CACCATCACCACAGCCTATT-3' | 5'-GCATTGTCCCACGAGTAAGT-3' |
| Rab3d | 5'-GTGTAGAAACGGAAGTGGAGAA -3' | 5'-CAAGAGTCCTCATGTGGAGAAG -3' |

| | | |
|---|---|---|
| Rab43 | 5'-CACACCATGAGGGCTGTATT-3' | 5'-GTTCTGTTCCACTCCAGGTTAG-3' |
| Rab4b | 5'-CCTACCTGAAGAACTCCCAAAG-3' | 5'-GAACAGGCCTCCAGGTAATAAA-3' |
| Rab5b | 5'-CAATGACAGGGCAGCTAGAA-3' | 5'-CAGACACCCTCAAACACCTAATA-3' |
| Rab5c | 5'-CAATGACCCGACTGGAATCTAC-3' | 5'-CCGGCCTAGGATCAAAGTTATG-3' |
| Rab7b | 5'-TGTCTTCACACTGCACAGATAG-3' | 5'-GTGGAAGGCATGAGGTATGAA-3' |
| Rel B | 5'-CTTCCAGCTTCCTCATCCTG-3' | 5'-CCTCTTCGGACTCAGCATTC-3' |
| RXRa | 5'-AGCCATTGTCCTGTTCAACC-3' | 5'-CCTAGGTGGCTTGATGTGGT-3' |
| SHP/Nr0b2 | 5'-CTGGTTGAGCGCCTGAGAC-3' | 5'-CTGCCTGGATGCCCTTTATC-3' |
| TCP1 | 5'-GCCTTGGGTGTCTCACATTA-3' | 5'-CAGTGCAACACTAAGCAGAAAG-3' |
| TRAP1 = HspC5=HSP75 | 5'-CCAGTGATGCCTTGGAGAAA-3' | 5'-GCCAGTGTCCTGAATGGTAATA-3' |
| XBP spliced | 5'-GAGTCCGCAGCAGGTG-3' | 5'-GTGTCAGAGTCCATGGGA-3' |
| XBP uncut | 5'-GAATGGACACGCTGGATCCT-3' | 5'-GCCACCAGCCTTACTCCACTC-3' |

**Supplementary Table 2**: List of antibodies-

**Supplementary Table 2. Antibodies used in immunoassays and IFC**

| **Antibody/Species** | **Source/Catalog Number/Dilution** |
|---|---|
| Actin/Mouse | Cell signaling Technology/#3700/1:4000 |
| AMPK/Rabbit | Cell signaling Technology/#2532/1:1000 |
| ATF6/Rabbit | Santa Cruz/sc-22799/1:500 |
| BIP/Rabbit | Cell signaling Technology/#3183/1:1000 |
| CHOPP/Rabbit | Cell signaling Technology/#2895/1:1000 |
| K18/TROMA-I/Rat | Developmental Studies Hybridoma Bank/ AB_531826/1:500 |
| K19/Rat | Developmental Studies Hybridoma Bank/1DB-001-0000868971/1:200 |
| K8/TROMA-I/Rat | Developmental Studies Hybridoma Bank/ AB_531826/1:500 |
| elF2α/Rabbit | Cell signaling Technology/#9722/1:1000 |
| FXR/Mouse | R&D Systems/PP-A9033A-00/1:1000 |
| Gadd34/Goat | Santa Cruz/sc-8832/1:500 |
| GAPDH/Mouse | Novus Bilogicals/NB300-21/1:3000 |
| HSF1/Rabbit | Cell signaling Technology/#4356/1:1000 |
| HSP70/Rabbit | Cell signaling Technology/#4872/1:1000 |
| HSP90/Rabbit | Cell signaling Technology/#4874/1:1000 |
| IRE1a/Rabbit | Santa Cruz/sc-20790/1:500 |
| LC3/Rabbit | MBL/PM036/1:1000 |
| NF-κB/p65/Rabbit | Cell signaling Technology/#8242/1:1000 |
| Nqo1/Rabbit | Abcam/ab34173/1:3000 |
| P62/SQSTM1/Mouse | Abnova/H00008878-M01/1:1000 |
| Phospho-4E-BP1/Rabbit | Cell Signaling Technology/#9459/1:1000 |
| Rab11/Mouse | BD Pharmigen/610656/1:1000 |
| Rab5/Mouse | BD Transduction/610724/1:1000 |
| Rab7/Mouse | Sigma/R8779/1:1000 |
| Rabenosyn 5/Goat | Abcam/ab21196/1:1000 |
| Rabex-5/Mouse | BD Transduction/612558/1:1000 |
| Spartin/Goat | Santa Cruz/sc-49521/1:500 |
| Total-4E-BP1/Rabbit | Cell Signaling Technology/#9452/1:1000 |
| XBP/Rabbit | Santa Cruz/sc-7160/1:500 |
| Alexa-488-labeled anti-Rabbit 2nd Ab/Goat | InVitrogen/A-11034/1:500 |
| Cy3-labeled anti-Rat 2nd Ab/Donkey | Jackson ImmunoResearch Laboratories Inc/712-165-150/1:500 |

| | |
|---|---|
| HRP-labeled anti-Goat 2nd Ab /Donkey | Jackson ImmunoResearch Laboratories Inc/705-035-147/1:5000 |
| HRP-labeled anti-Mouse 2nd Ab/Goat | Jackson ImmunoResearch Laboratories Inc/115-035-062/1:5000 |
| HRP-labeled anti-Rabbit 2nd Ab/Goat | Jackson ImmunoResearch Laboratories Inc/111-035-045/1:5000 |
